# Transfer entropy predicts pupillary response and cognitive effort during a tracking task

**DOI:** 10.1101/2025.04.01.646647

**Authors:** Simon Boylan, Sze Ying Lam, Alexandre Zénon

## Abstract

Cognitive cost plays a crucial role in determining action choices and task engagement. We have proposed that cognitive cost relates to the amount of Bayesian inference required, quantified using relative entropy. In continuous tasks, such as visuomotor tracking, this demand can be estimated using transfer entropy (TE), which accounts for the information transferred from input to output variables. This study aimed to test experimentally the relationship between TE and markers of cognitive cost.

We designed a continuous tracking task, manipulating target speed, predictability, and response lag. The effects of task variables on performance, subjective effort and pupil-linked arousal were assessed using Bayesian mixed models. We disentangled the effect of TE from that of other correlated variables (predictability, acceleration, spatial error), by using model comparison and mediation models. Results showed that TE was mostly affected by target predictability and impacted significantly on subjective effort and pupil response. While the variations of pupillary response induced by task conditions were accounted for by TE only, subjective effort was only partially predicted by TE, and depended also on target predictability.

These findings suggest that the cognitive demand of motor control tasks can be quantified using TE, which relates to both subjective effort and pupillary response. However, the multifaceted nature of effort in motor tasks and the limitations of subjective measures highlight the need for more nuanced tools to isolate mental and physical effort.

## 2. Introduction

Why is it more appealing to browse on internet, or read a book, rather than do academic work? Decades of research have proposed different motivation-based models that explain action choices and energy put into task (1–3). A consensus is emerging around trade-off models of cognitive control, which emphasize a balance between rewards and costs (4–6). The cost of cognition manifests as the aversiveness of cognitive tasks, often accompanied by an unpleasant feeling of exertion, referred to as cognitive effort (7).

Cognitive cost influences the choice of which action to undertake, as well as its intensity in terms of cognitive control (6,8). Despite its central role, the physiological origin of the cost of cognition is still not established (5). What seems clear however, is that cognitive task exertion is not linked to increased energy consumption, since the amount of energy consumed by the brain is the same whether at rest or doing tasks (9,10).

Inspired by decades of research that have used information theory as a framework to study neural processing (11–23), and by the Bayesian approach to brain function (24), we have proposed that the cognitive demand of a task relates to how much Bayesian inference it requires, and can be quantified in terms of relative entropy: the (Kullback-Leibler) divergence between the prior and posterior beliefs over task variables (25). Interestingly, the pupillary response to cognitive processing has been shown to relate to the same relative entropy quantity, and would thus provide us with an objective, specific measure of task demand (26).

We have also proposed that in the context of motor control, and in particular in visuomotor tracking tasks, such information-theoretic measures of cognitive task demand could be estimated by means of the transfer entropy (TE), i.e. the amount of information transferred from input to output variables (27). The total information shared by inputs and outputs can be decomposed into a feedforward, predictive component and a feedback, error-correcting component. TE accounts for the feedback component of tracking, which represents the amount of information needed to update the control output given novel sensory data. According to our theory, this TE estimate should translate into computational cost and subjective effort.

In the present study, we designed a continuous tracking task to explore the relationship between information processing, pupil size and cognitive demand in a motor control context. We predicted that subjective effort and pupillary response to the task should relate specifically to TE. We manipulated the speed and predictability of the target trajectory on the screen, as well as the lag between subjects’ tracking response and its effect on the screen cursor. This response lag manipulation was added in order to perturb prediction mechanisms without changing the nature of the signal. We assessed the effect of these variables on performance, subjective effort (measured through NASA-TLX questionnaires) and pupil size, and investigated the mediating influence of TE on these effects.

## 3. Materials and Methods

### Participants

Twenty-four right-handed subjects (7 males) between 18 and 29 years old were included in this study. All subjects had normal or corrected-to-normal vision, and declared no history of neurological or mental disorders. They signed an informed consent form before participating in the study and were financially compensated for their participation. The protocol was approved by the local Ethics Committee and adhered to the principles expressed in the Declaration of Helsinki.

### Experimental design and procedure

Participants were seated in front of an OLED computer screen (Asus 27” OLED - ROG Swift PG27AQDM; 60×34 cm, resolution: 2560×1440) and were holding a joystick in their right hand. The joystick had its central spring removed in order to keep constant resistance throughout its course. The instruction given to participants in this visuo-motor tracking task was to aim at placing the joystick-controlled cursor (2°-wide blue square) on the moving target (white, 1°-wide vertical bar) as best as they could.

Throughout the whole experiment, subjects had to keep their head still on the head-stand anchored to the desk. They were also instructed to fixate the crosshair at the center of the screen during the trials. Both measures were taken to minimize head and eye movements that could compromise pupil diameter recordings.

Three task properties were manipulated in this experiment: signal predictability, signal speed, and added motor delay (AMD). The tracking signals in this experiment were generated by processing white noise through a sinusoidal filter: *a*_1_ *x*_*t*_=*θ*_*t*_ +*a*_2_ *x*_*t−*1_+ *x*_*t−*2_, with 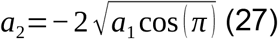.

We generated signals of different levels of complexity using *a*_1_ parameter values ranging from 1 to 1.08. Then, the signals of each complexity level were either slowed down (upsampled) or sped up (downsampled). The level of predictability of the signals was quantified by means of sample entropy (28). Among all the signals generated, we selected 5 speed-entropy conditions that were approximately orthogonal (correlation coefficient =-0.06; see Figure 1). On the contrary, the acceleration of the target was found to relate to both Target Speed (correlation coefficient=0.71) and sample entropy, called henceforth Target Unpredictability (0.57), such that the variance explained by a linear model of the form *acceleration ∼ speed + unpredictability* was 87%. Each of the 5 signal conditions were tested in 2 different motor delay conditions: 0ms or 280ms. There were therefore in total 5 (groups) x 2 (AMD) = 10 different conditions in this experiment. Participants executed 6 blocks of 10 trials each, in pseudorandom order, for a total of 60 trials (6 per condition). Each trial lasted 25 seconds.

**Figure 1.**
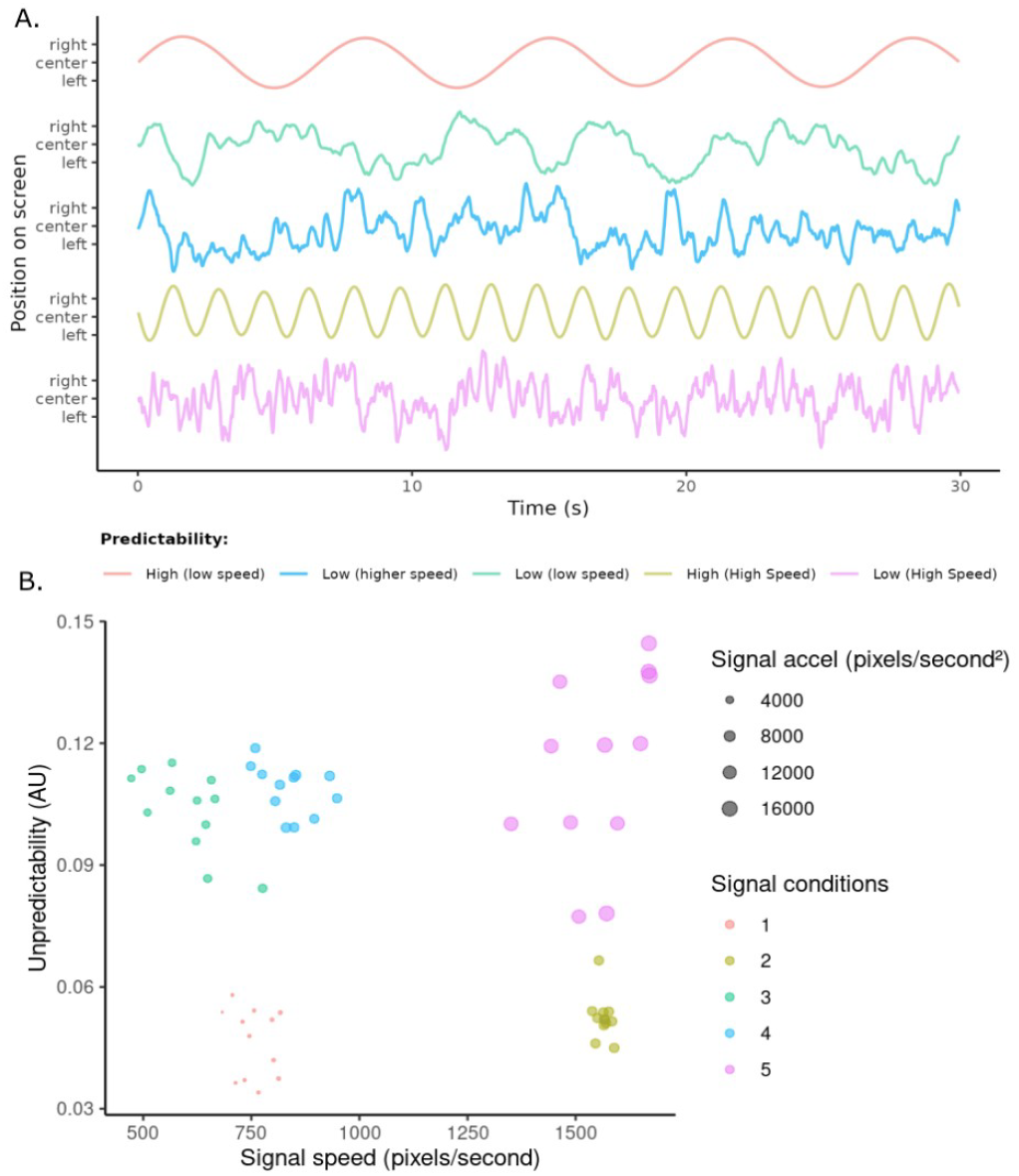
A. Example time courses of the five signal conditions used in the experiment. B. Average speed, unpredictability (sample entropy) and acceleration of each signals used in the experiment.

In three trials of every condition, selected pseudorandomly, subjects were presented with two selected questions from the NASA Task Load indeX questionnaire (Hart and Staveland, 1988) on perceived level of mental demand, physical demand and effort associated with the trial they had just performed (see explicit questions in Supplementary Table 1). The complete questionnaire was first shown to the subjects before the experiment so as to ensure their understanding of the subtle differences between the different items, including the differences between mental demand, physical demand, effort and performance. After the target and cursor both disappeared from the screen at the end of the trial, the first question was presented with a ruler scale at the bottom of the screen. Subjects could then respond to the question by placing the cursor on the scale using the joystick and then validate the chosen location with a key press.

### Data and statistical analyses

The feedback component was computed as a TE term relating visual input to motor output, in accordance with our previous publication (27) and defined as the mutual information between the output *Y* at time *t* and the input *X* at times *t-VMD* and *t-VMD-1*, conditioned on *Y* at times *t-1, t-VMD* and *t-VMD-1: TE*=I(*Yt*;{*Xt−VMD,Xt−VMD−1*}|{*Yt−VMD,Yt−VMD−1,Yt−1*}), with VMD the visuomotor delay. We used the Kraskov method for estimating conditional mutual information (29). The corresponding code can be found on Github (https://github.com/alexandre-zenon/TrackingInfoRate).

In the pupil data, blink epochs were isolated and replaced with linear interpolation. The signal was then downsampled to 10 Hz and filtered with a non-causal Chebyshev filter with a 5-order 0.1Hz cutoff frequency (chosen so as to ensure 40dB attenuation and less than 1dB ripple within the bandpass). The filtered pupil signal was then averaged over the trial duration, excluding the first 10 seconds, allowing for proper stabilization of the signal.

All statistical analyses were Bayesian mixed models (30) run with the package brms (31) in the R language (version 4.4.1). One of the main advantages of Bayesian statistics is the ability to account precisely for prior knowledge through the use of prior distributions. We chose weakly informative prior distributions for all models, based on normal distributions for predictor parameters and Student’s t distributions for intercept and standard deviation parameters. We performed prior predictive checks by looking at the output of the models with all parameters distributed according to the prior and comparing the distribution of the sampled data with the actual data (32). In general, since all our variables were z-scored prior to being used in the model, we chose normal distributions with 0 mean and SD=1 for all population-level effects (mildly informative), whereas priors for the intercept and standard deviation parameters were distributed according to Student’s t distributions, with v=10, μ=0, and σ = 1 (weakly informative).

We ran 4 independent Markov Chain Monte-Carlo (MCMC) for each model, with 10,000 iterations and a warmup of 5,000 iterations. We reported 95% highest density intervals (HDI), a Bayesian equivalent to confidence intervals (30). We assessed the convergence of the chains by examining Effective Sample Size (ESS) and the Gelman-Rubin statistic 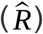. R-squared values were obtained with the method described by Gelman and colleagues (33). Model comparisons were performed by comparing expected log predictive densities (ELPD), estimated by means of Leave-One-Out (LOO) cross-validation (34).

## 4. Results

### Transfer Entropy

We compared different models to account for Transfer Entropy (TE) data: Target Speed, Target Unpredictability, Target Acceleration, or signal condition (5 groups of speed-entropy-acceleration combinations, see Figure 1). All models either included motor delay, or not, resulting in a total of 6 models. We found that the best model in terms of expected log predictive density (ELPD), estimated by means of Leave-One-Out (LOO) cross-validation, was the one with signal condition and motor delay as predictors (R^2^=0.87±0.002; ELPD difference with next best model= 87±15 [estimate±SE]; see Supplementary Figure 1). Models are considered to be statistically different if their ELPD difference is larger than 1.96*standard error. We nevertheless examined the result of the model that included Target Speed, Target Unpredictability and Motor Delay as predictors, in order to get a feeling of the relationship between those variables and TE (formula: TE *∼ Motor_Delay * Speed * Unpredictability + (Motor_Delay * Speed * Unpredictability* | *subject)*.

We found that all variables had a significant positive impact on TE, but as expected, target Unpredictability had the strongest effect by far (see Fig. 2 and Table S2). This confirms our hypothesis that more unpredictable (higher sample entropy), jerky (higher acceleration) and swift (higher speed) targets require more visual information processing to reduce tracking error. Similarly, compensation of the Motor Delay needs extra cognitive control, which translates also to higher TE.

**Figure 2.**
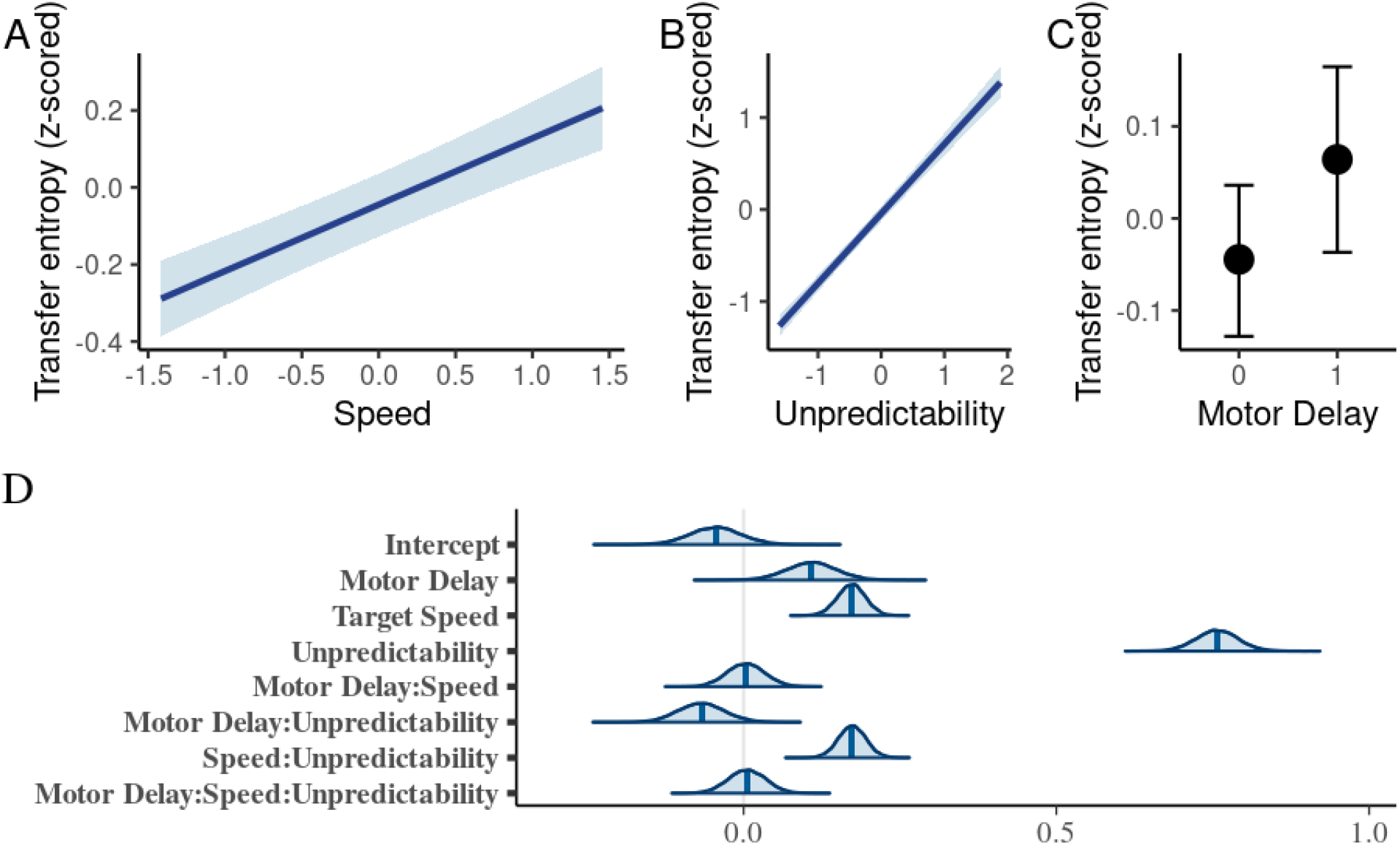
Effect of Target Speed, Target Unpredictability and Motor Delay on Transfer Entropy. A. Effect of Target Speed (z-scored) on Transfer Entropy. B. Effect of Target Unpredictability (sample entropy, z-scored) on Transfer Entropy. C. Effect of Motor Delay on Transfer Entropy. D. Posterior distributions of predictor parameters. Target Unpredictability had the strongest impact on Transfer Entropy.

In the following analyses, we set out to determine the influence of TE on the other variables of interest (RMSE, NASA-TLX scores and pupillary responses) by including it as a predictor in the models.

### RMSE

We ran multiple models on the RMSE data to verify which predictor configuration (with either Signal Conditions, Acceleration, Speed, Unpredictability and TE) provided the best prediction of spatial error. Even though the model including TE as predictor highlighted a significant effect of TE on RMSE (HDI: 0.46-0.57), model comparisons showed that the model with Signal Condition and Motor Delay as predictors was the best model by far (see Figure 3 and Table S3; R2=0.82±0.004; EPLD difference with next best model= 143±27; formula: *spatial_error ∼ Motor_Delay * Signal_Condition + (Motor_Delay * Signal_Condition* | *subject))*.

**Figure 3.**
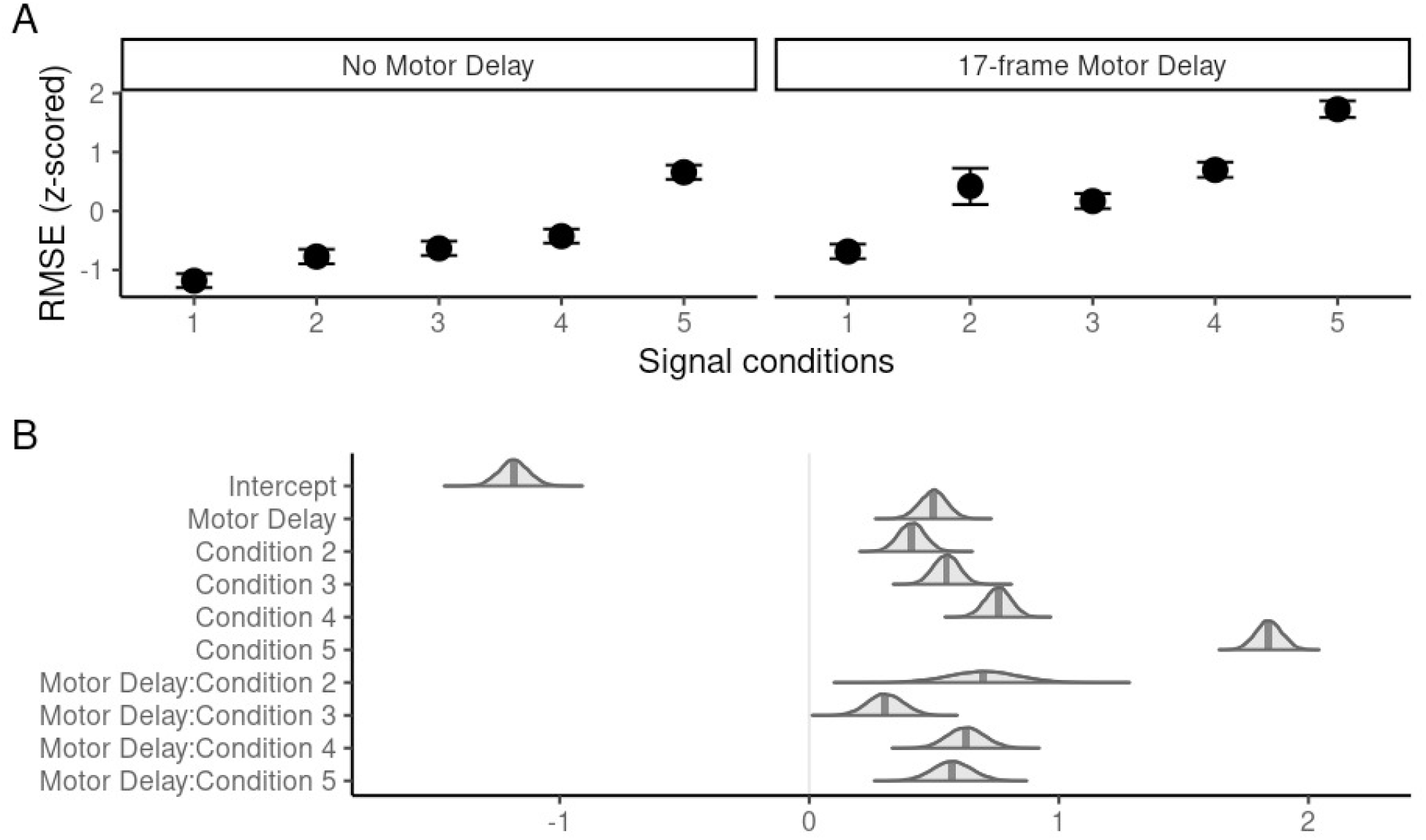
A. Effect of signal conditions and motor delay on RMSE. B. Distributions of predictor parameters.

The results from this model show that signal conditions that correspond to less predictable and faster target movements are harder to track, leading to more spatial error than with easier targets. However, this effect is not easily accounted for by simpler models that separate Target Unpredictability and Target Speed. Motor delay also leads to larger spatial errors overall.

### NASA-TLX

The three scores of the NASA questionnaire data correlated strongly with each other (coefficients from 0.48 to 0.84, see Table S4). We thus decided to run a principal component analysis to reduce the number of scores and highlight specific features. We found that the first principal component of the NASA scores (74% of the variance) corresponded to an average of the three scores, while the second principal component (21%) seemed to highlight what was specific to mental effort. The third component had negligible contribution to the variance (5%) (see Tables S5 & S6). Therefore, we performed a similar model comparison as above, using the two first NASA principal components as dependent variables. We found that the second principal component had no relationship to any of the other variables (best model had no significant effect). We thus only kept the first principal component corresponding to the average of all three scores, for the rest of the analyses.

Model comparison on the first principal component showed that the best models, with no statistically significant differences between them, were the ones with Motor Delay, Target Speed and Target Unpredictability (without speed-unpredictability interaction, R2=0.67±0.02; formula of model 1: *nasaPCA1 ∼ Motor_Delay * (Speed + Unpredictability) + (Motor_Delay * (Speed + Unpredictability)* | *subject)*), and the one with only Motor Delay and TE (0.63±0.02) as predictor variables (formula of model 2: nasaPCA1 ∼ Motor_Delay * TE + (Motor_Delay * TE | subject)).

Results from the first model revealed the stronger effect of Target Unpredictability on NASA-TLX’s first principal component with respect to the other variables (see see Figure 4A-C and Tables S7). The main effects of Target Speed and Motor Delay were not significant. Interestingly, we found a significant interaction between Speed and Motor Delay suggesting that participants only found the compensation of Motor Delay effortful when the target was faster.

**Figure 4.**
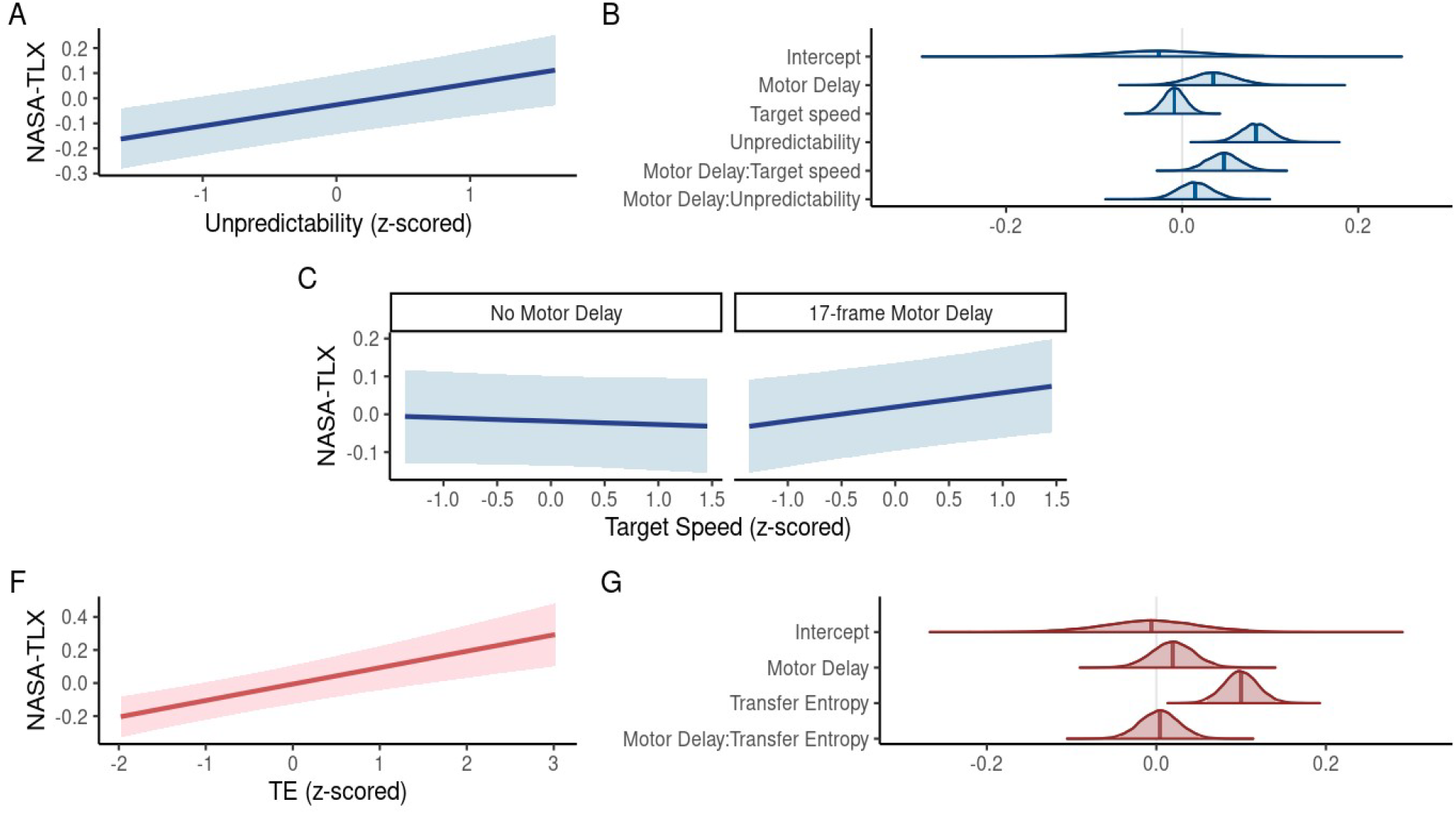
Results of the Speed-Unpredictability and Transfer Entropy models on the NASA-TLX scores. Difference in colors illustrate the different models. A. Effect of Target Unpredictability (sample entropy) on NASA scores. B. Posterior distributions of the parameters associated with the predictor variables in the Speed-Unpredictability model. C. Effect of Target Speed on the NASA scores as a function of Motor Delay. D. Effect of Transfer Entropy on NASA-TLX. E. Distribution of posterior parameters in the Transfer Entropy model.

The second model highlighted the strong effect of TE on subjective effort (see Figure 4F-G and Tables S8), confirming our hypothesis that TE could be interpreted as a measure of cognitive cost.

We then attempted to assess whether the TE variable mediated the effect of Target Unpredictability on the NASA-TLX score, as we would expect if TE was really the decisive feature that mattered for subjective effort. Our results showed an indirect effect of Target Unpredictability on the NASA-TLX score mediated by TE which was only at the edge of significance (0.034 [-0.001 0.069]) and which represented 37.87% of the total effect (see Table S9), thus only partially confirming our hypothesis.

### Pupil data

Model comparison showed that the best model accounting for pupillary responses was the one with TE as predictor (formula: *pupil_response ∼ TE + (TE* | *subject)*, followed by the one with target acceleration as predictor (ΔELPD=5.4±8.23, formula: *pupil_response∼ acceleration + (acceleration* | *subject)*). Other models provided clearly poorer fit to the data (next best model with Target Speed and Target Unpredictability: ΔELPD=11.9±10.39). Both the TE and Target Acceleration models showed very significant effects of their predictors on pupillary dilation (see Fig. 5 and Tables S10 and S11).

**Figure 5:**
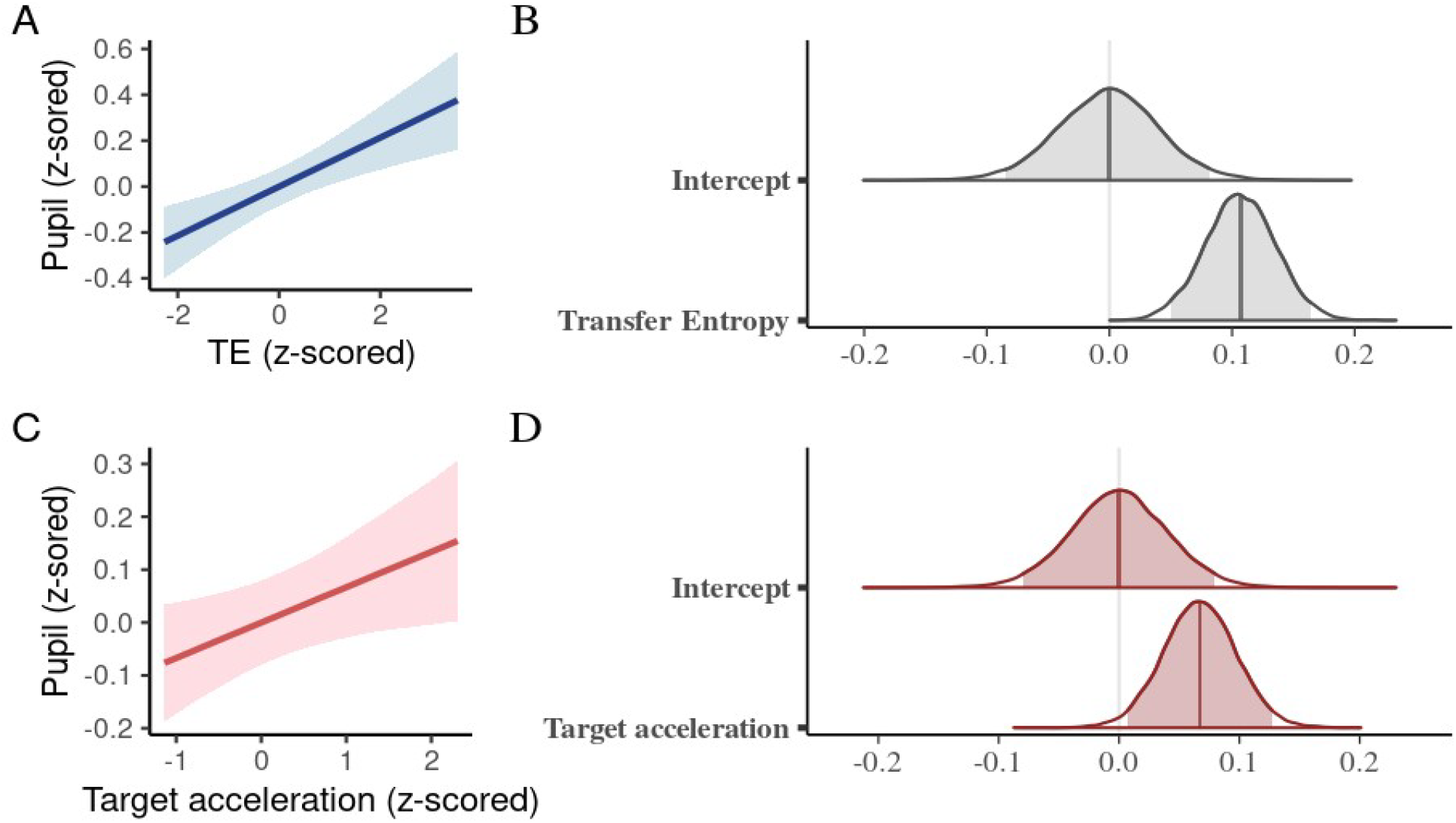
Results of the two best models of pupil data. Different colors are used to illustrate the two models. A. Effect of Transfer Entropy on pupil response. B. Distribution of posteriors of the predictor parameters for the model with Transfer Entropy. C. Effect of Target Acceleration on pupil response. D. Distribution of posteriors of the predictor parameters for the model with Target Acceleration.

Mediation analysis showed that the effect of Target Acceleration on pupil responses was fully mediated by the effect of TE (proportion mediated = 126%; indirect effect=0.088 [0.005 0.180]; direct effect=-0.019 [-0.134 0.093], see Table 12).

These results show that pupil response was strongly affected by TE and that the effect of acceleration was fully mediated by its indirect effect on TE, again confirming our view that TE is the fundamental driver of pupillary dilation

## 5. Discussion

This study investigated the relationship between cognitive effort, transfer entropy (TE) and pupillary response during a continuous visuomotor tracking task. By manipulating target speed, unpredictability (sample entropy), and motor delay, we aimed to assess how these factors influenced subjective (NASA-TLX scores) and objective (pupil size) measures of cognitive effort. Our findings provide valuable insights into how information processing, task difficulty, and physiological responses are interrelated, though they also highlight several complexities that warrant further exploration.

The results partially supported our hypothesis that TE could serve as a key measure of cognitive effort. Specifically, TE was found to be a strong predictor of pupillary response and accounted for the effects of all other variables, confirming that it captures the cognitive demands imposed by the task. However, the effect of TE on subjective effort was a bit less clear, with a marginally significant mediation of the relationship between target unpredictability and subjective effort, suggesting that while it reflects information processing requirements, it may not be the sole factor influencing perceived effort.

This discrepancy between objective and subjective measures likely stems from the multifaceted nature of effort in a motor task. While TE successfully captures the cognitive component related to processing sensory feedback and adjusting motor output, subjective effort ratings were probably influenced by both physical and mental aspects of the task, complicating the interpretation of TE’s role. This finding aligns with prior research indicating that subjective assessments of cognitive effort are often influenced by a blend of mental workload and physical demands, particularly in tasks involving motor control (35). The NASA-TLX questionnaire, though widely used to measure subjective effort, revealed some limitations in this study. The three subscales—mental demand, physical demand, and overall effort—were highly correlated, suggesting that participants had difficulty distinguishing between the different components of effort in a motor task. This overlap likely explains why TE was only a partial predictor of subjective effort, as the questionnaire may not have effectively isolated the mental aspects of cognitive load from the physical effort involved in controlling the joystick. This points to the need for more nuanced tools to measure mental and physical effort separately, especially in tasks where both are heavily intertwined.

One of the most robust findings in our study was the strong relationship between TE and pupillary dilation. Pupillary responses have long been associated with cognitive load (36), and our results confirm that pupil size increases with higher task demands, as quantified by TE (26,27). Interestingly, while target speed and acceleration were found to have significant effects on task performance and effort, their influence on pupil size was secondary to that of TE. This reinforces the idea that cognitive effort, as measured by TE, plays a primary role in driving physiological responses to task demands. This highlights the potential of using TE as a reliable, objective measure of cognitive effort, especially in contexts where pupil size can be easily monitored.

The findings of this study suggest that TE could serve as a generalizable measure of cognitive effort beyond visuomotor tracking tasks. TE quantifies the flow of information between input and output variables, making it a potentially valuable tool for studying cognitive load in diverse domains such as decision-making, problem-solving, and language processing. For example, in complex decision-making scenarios, TE could capture the degree of uncertainty resolution, aligning with theories that link cognitive effort to Bayesian inference and information gain (25). Similarly, in linguistic processing, where predictive mechanisms play a central role, TE could offer insights into how individuals manage syntactic and semantic complexity (37,38). Expanding TE-based analysis to these domains could provide a unified, quantitative approach to understanding cognitive effort across disciplines, enhancing models of cognitive control and workload (4). Future research should investigate whether TE can reliably predict cognitive load across various modalities and how it compares to traditional measures such as reaction times, pupil dilation, and neural activity patterns.

## Acknowledgments

The authors wish to thank Oleg Solopchuk for his helpful comments on the manuscript.

## Ethical Statement

The study was approved by the Ethical Review Board and adhered to the principles of the Declaration of Helsinki. All participants were financially compensated for their participation (10€ per hour) and provided written informed consent to participate.

## Data Accessibility

*Data processing scripts and datasets can be found on Github (https://github.com/alexandre-zenon/TrackingInfoRate)*.

## Authors’ Contributions

Conceptualization: AZ (lead), SYL (equal)

Data curation: AZ(lead), SB(supporting)

Formal analysis: AZ (lead), SB (supporting)

Funding acquisition: AZ

Project administration: AZ (lead), SB (equal)

Visualization: AZ(lead), SB (supporting)

Writing – original draft: AZ (lead), SB (equal)

Writing – review & editing: SB (lead), AZ (equal), SYL (supporting)

