## Supplemental material for "Transfer entropy predicts pupillary response and cognitive effort during a tracking task"

### **Supplementary Material**

#### Supplementary tables

Table S1: NASA-TLX questionnaires items on task demand and effort

| Quelle a été l’importance de l’activité mentale et intellectuelle requise (ex. réflexion, décision, calcul, mémorisation, observation, recherche etc.)?  La tâche vous a-t-elle parue simple, nécessitant peu d’attention (faible) ou complexe, nécessitant beaucoup d’attention (élevée)? | How much mental and perceptual activity, was required (eg., thinking, deciding, calculating, remembering, looking, searching, etc)?  Was the task easy or demanding, simple or complex, exacting or forgiving? |
| --- | --- |
| Quelle a été l’importance de l’activité physique requise (ex. pousser, porter, tourner, marcher, activer, etc.) ?  La tâche vous a-t-elle paru facile, peu fatigante, calme (faible) ou pénible, fatigante, active (élevée)? | How much physical activity was required (e.g., pushing, pulling, turning, controlling, activating, etc.)?  Was the task easy or demanding, slow or brisk, slack or strenuous restful or laborious? |
| Quel degré d’effort avez-vous dû fournir pour exécuter la tâche demandée, (mentalement et physiquement)? | How hard did you have to work (men-  tally and physically) to accomplish  your level of performance? |

Table S2 : Transfer Entropy Bayesian linear model’s summary.

|  | Estimate | Est.Error | CI.Lower | CI.Upper | Rhat | Bulk_ESS | Tail_ESS |
| --- | --- | --- | --- | --- | --- | --- | --- |
| Intercept | -0.04 | 0.04 | -0.13 | 0.04 | 1 | 8388.36 | 11657.36 |
| Motor Delay | 0.11 | 0.04 | 0.03 | 0.19 | 1 | 15678.07 | 15480.42 |
| Speed | 0.17 | 0.02 | 0.13 | 0.22 | 1 | 22206.28 | 14927.33 |
| Unpredictability | 0.76 | 0.03 | 0.69 | 0.83 | 1 | 12459.5 | 14887.19 |
| Motor Delay:Speed | 0 | 0.03 | -0.05 | 0.06 | 1 | 24770.55 | 15480.86 |
| Motor Delay:Unpredictability | -0.07 | 0.04 | -0.14 | 0.01 | 1 | 20834.02 | 16245.55 |
| Speed:Unpredictability | 0.17 | 0.02 | 0.13 | 0.22 | 1 | 21134.89 | 15451.37 |
| Motor Delay:Speed:Unpredictability | 0.01 | 0.03 | -0.05 | 0.06 | 1 | 28353.62 | 16124.99 |

Table S3: Spatial error (RMSE) Bayesian linear model’s summary.

|  | Estimate | Est.Error | CI.Lower | CI.Upper | Rhat | Bulk_ESS | Tail_ESS |
| --- | --- | --- | --- | --- | --- | --- | --- |
| Intercept | -1.18 | 0.06 | -1.3 | -1.06 | 1 | 4253.55 | 7333.52 |
| Motor Delay | 0.5 | 0.05 | 0.39 | 0.6 | 1 | 7137.76 | 11415.03 |
| Condition 2 | 0.41 | 0.05 | 0.31 | 0.51 | 1 | 9461.69 | 12734.16 |
| Condition 3 | 0.55 | 0.05 | 0.45 | 0.65 | 1 | 9306.75 | 12681.59 |
| Condition 4 | 0.76 | 0.05 | 0.65 | 0.86 | 1 | 9198.23 | 13092.27 |
| Condition 5 | 1.84 | 0.05 | 1.74 | 1.94 | 1 | 9348.01 | 12411.16 |
| Motor Delay:condition 2 | 0.7 | 0.14 | 0.42 | 0.97 | 1 | 7398.42 | 10791.25 |
| Motor Delay:condition 3 | 0.31 | 0.07 | 0.17 | 0.45 | 1 | 8861.09 | 11643.37 |
| Motor Delay:condition 4 | 0.63 | 0.07 | 0.48 | 0.78 | 1 | 8259.38 | 12994.99 |
| Motor Delay:condition 5 | 0.57 | 0.08 | 0.42 | 0.73 | 1 | 8350.76 | 11760.79 |

Table S4: Correlation matrix of each NASA-TLX item responses.

|  | NASA 1 | NASA 2 | NASA 3 |
| --- | --- | --- | --- |
| NASA 1 | 1.00 | 0.47 | 0.84 |
| NASA 2 | 0.47 | 1.00 | 0.53 |
| NASA 3 | 0.84 | 0.53 | 1.00 |

Table S5: Importance of components of NASA-TLX results. First and second components have respectively 74.2 and 20.6 % while the third component only has 5.2 %.

| Importance of Components | nasaPC1 | nasaPC2 | nasaPC3 |
| --- | --- | --- | --- |
| Standard deviation | 0.34 | 0.18 | 0.09 |
| Proportion of Variance | 0.74 | 0.21 | 0.05 |
| Cumulative proportion | 0.34 | 0.18 | 0.09 |

Table S6: Decomposition of the three independent components of NASA-TLX. The first component is an average of the three scores, while the second component highlights what is specific to mental demand.

|  | nasaPC1 | nasaPC2 | nasaPC3 |
| --- | --- | --- | --- |
| NASA 1 | 0.60 | 0.42 | 0.68 |
| NASA 2 | 0.52 | -0.85 | 0.07 |
| NASA 3 | 0.61 | 0.31 | -0.73 |

Table S7: Effort (NASA Princip*al component*) Bayesian linear first model’s summary.
*nasaPCA1 ~ Delay * (Speed + Unpredictability) + (Delay * (Speed + Unpredictability) | subject)*

|  | Estimate | Est.Error | CI.Lower | CI.Upper | Rhat | Bulk_ESS | Tail_ESS |
| --- | --- | --- | --- | --- | --- | --- | --- |
| Intercept | -0.03 | 0.06 | -0.14 | 0.09 | 1 | 2251.53 | 4525.39 |
| Motor Delay | 0.04 | 0.03 | -0.02 | 0.09 | 1 | 14483.15 | 14215.79 |
| Speed | -0.01 | 0.01 | -0.03 | 0.02 | 1 | 23425.64 | 15758.8 |
| Unpredictability | 0.08 | 0.02 | 0.05 | 0.12 | 1 | 11895.36 | 13370.77 |
| Motor Delay:Speed | 0.05 | 0.02 | 0.01 | 0.08 | 1 | 22111.29 | 15516.44 |
| Motor Delay:Unpredictability | 0.01 | 0.02 | -0.02 | 0.05 | 1 | 24077.81 | 15067.2 |

Table S8 : Effort (NASA Princip*al component*) Bayesian linear second model’s summary.
*nasaPCA1 ~ Delay * TE + (Delay *TE | subject)*

|  | Estimate | Est.Error | CI.Lower | CI.Upper | Rhat | Bulk_ESS | Tail_ESS |
| --- | --- | --- | --- | --- | --- | --- | --- |
| Intercept | -0.01 | 0.06 | -0.13 | 0.11 | 1 | 2543.06 | 4596.65 |
| Motor Delay | 0.02 | 0.03 | -0.03 | 0.07 | 1 | 17461.04 | 14175.98 |
| Transfer Entropy | 0.1 | 0.02 | 0.06 | 0.14 | 1 | 10392.63 | 12338.15 |
| Motor Delay:Transfer Entropy | 0 | 0.02 | -0.04 | 0.05 | 1 | 19504.02 | 14523.18 |

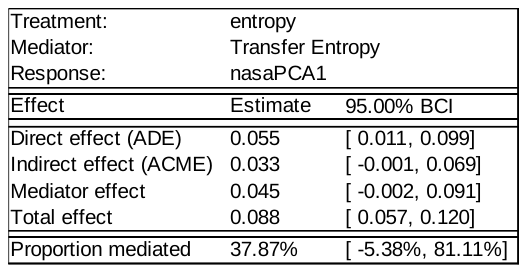
Table S10: Pupil response Bayesian linear model’s summary.
P*upil_response ~ target_acceleration + (target_acceleration | subject)*

Table S9: Transfer Entropy mediates 37.87 % of Target Unpredictability's effect
 on NASA-TLX’s principal component.

|  | Estimate | Est.Error | CI.Lower | CI.upper | Rhat | Bulk_ESS | Tail_ESS |
| --- | --- | --- | --- | --- | --- | --- | --- |
| Intercept | 0 | 0.04 | -0.08 | 0.08 | 1 | 10640.04 | 13167.53 |
| Target Acceleration | 0.07 | 0.03 | 0.01 | 0.13 | 1 | 18383.91 | 13528.72 |

Table S11 : Pupil response Bayesian linear model’s summary.
P*upil_response ~ transfer_entropy + (transfer_entropy | subject)*

|  | Estimate | Est.Error | l-95% CI | u-95% CI | Rhat | Bulk_ESS | Tail_ESS |
| --- | --- | --- | --- | --- | --- | --- | --- |
| Intercept | 0 | 0.04 | -0.08 | 0.08 | 1 | 13000.22 | 13541.55 |
| Transfer Entropy | 0.11 | 0.03 | 0.05 | 0.16 | 1 | 26786.24 | 14466.5 |

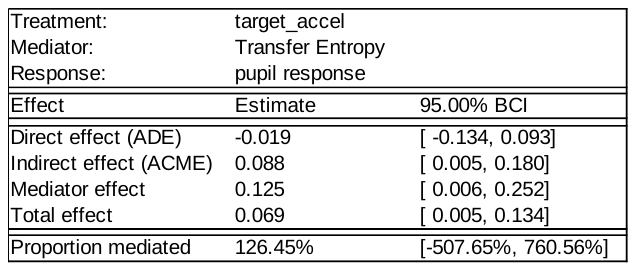

Table S12: Transfer Entropy fully mediates the effect of acceleration on pupil response. Previous significant direct effect isn’t statistically different from 0 ([ -0.134, 0.093]) when including Acceleration’s mediation.

#### Supplementary Figures

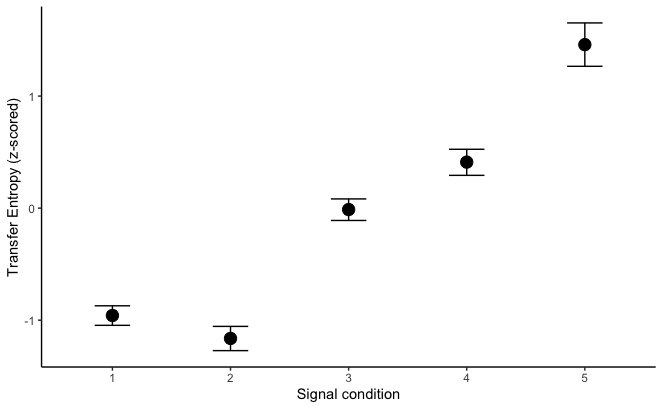
Figure S1 : Transfer Entropy as a function of signal conditions. Error bars illustrate 95 % HDI.
